# The synaptic organization in the *C. elegans* neural network suggests significant local compartmentalized computations

**DOI:** 10.1101/2021.12.30.474568

**Authors:** Rotem Ruach, Nir Ratner, Scott W. Emmons, Alon Zaslaver

## Abstract

Neurons are characterized by elaborate tree-like dendritic structures that support local computations by integrating multiple inputs from upstream presynaptic neurons. It is less clear if simple neurons, consisting of a few or even a single neurite, may perform local computations as well. To address this question, we focused on the compact neural network of *C. elegans* animals for which the full wiring diagram is available, including the coordinates of individual synapses. We find that the positions of the chemical synapses along the neurites are not randomly distributed, nor can they be explained by anatomical constraints. Instead, synapses tend to form clusters, an organization that supports local compartmentalized computations. In mutually-synapsing neurons, connections of opposite polarity cluster separately, suggesting that positive and negative feedback dynamics may be implemented in discrete compartmentalized regions along neurites. In triple-neuron circuits, the non-random synaptic organization may facilitate local functional roles, such as signal integration and coordinated activation of functionally-related downstream neurons. These clustered synaptic topologies emerge as a guiding principle in the network presumably to facilitate distinct parallel functions along a single neurite, effectively increasing the computational capacity of the neural network.

## Introduction

Neurons carry impressive computational repertoires. Much of these are due to the elaborate dendritic structures which simultaneously integrate and process multiple inputs ^1–6^. For example, songbirds implement a reliable coincidence detector based on non-linear summation of multiple inputs to the dendritic tree ^7^. Key to these computations is the relative position of the synapses along the dendritic tree ^8–10^.

In organisms whose neurons lack elaborate dendritic structures, the prospects for such local computations remain unclear. For example, the vast majority of the 302 neurons in *C. elegans* nematodes lack elaborate tree-like structures ^11^. In fact, many of these neurons consist of a single (unipolar) neurite extension ^12^, on which input and output synaptic sites are intermittently positioned (**Figure 1b**). An intriguing emerging question is whether local dendritic-like computations are possible in these simply-structured neurites.

**Figure 1.**
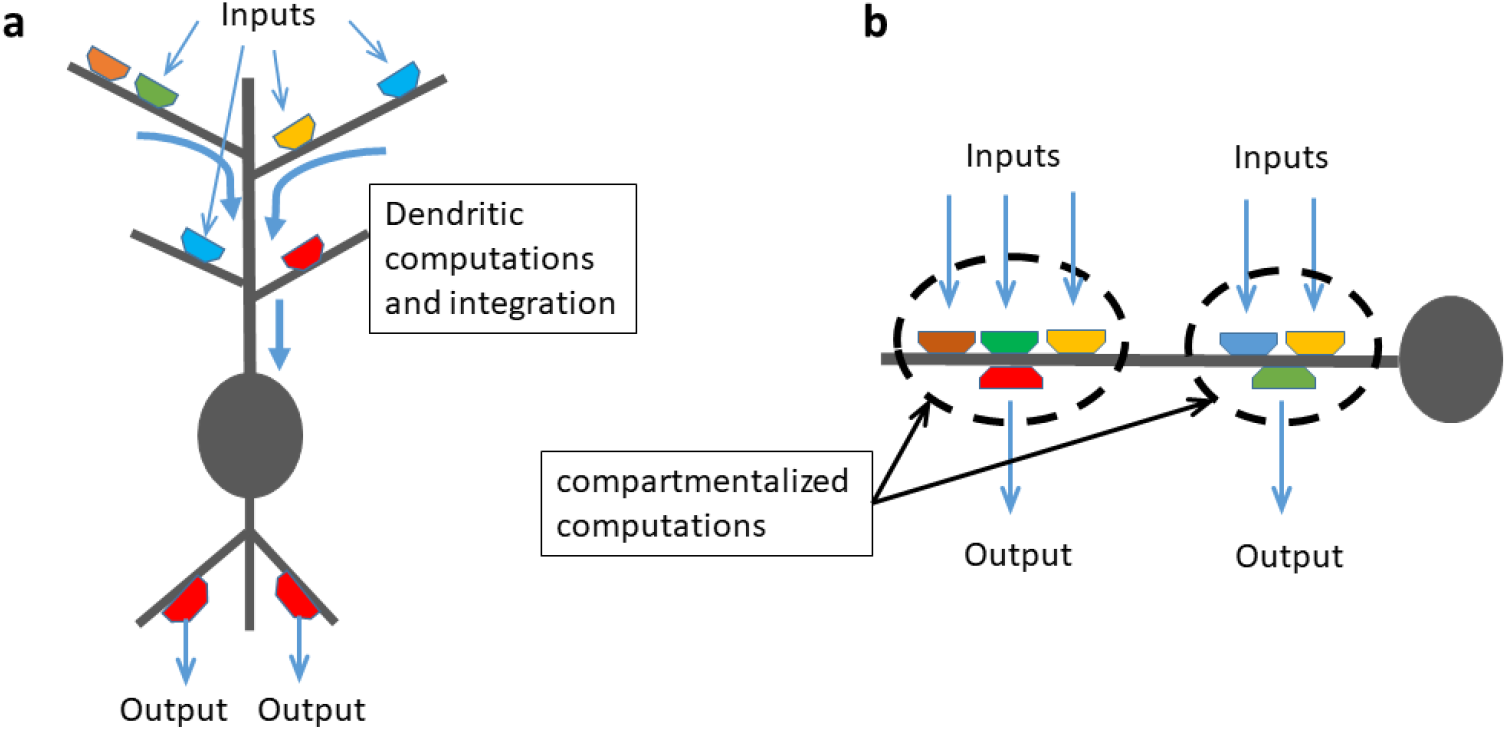
Dendritic computation in neurons with elaborate dendritic structure (a), and possible compartmentalized activity in discrete modules along neurites of simply-structured neurons (b). (a) A scheme of a typical arborized neuron with elaborate dendritic and axonal structures. Multiple inputs from presynaptic neurons are processed along the different branches of the dendrites. These signals are integrated and the output is transmitted via the axons to the downstream neurons. In this case, each dendrite serves as an intermediate integration site and the neuron operates as a single computational unit within the network. (b) A depiction of a simply-structured neuron consisting of a single neurite. Pre- and post-synaptic contacts are intermittently dispersed along the neurite. If these synaptic contacts form discrete clustered organizations, then each of these clusters can be viewed as a single compartmentalized module. Inputs from upstream neurons are locally integrated and the output to downstream neurons is transmitted via local synaptic contacts. In this case, several computations may be implemented on each neurite and their independent parallel function effectively increases the computational capacity of the neuron. Colored hexagons mark synaptic contacts and gray filled circles represent the cell soma.

In the absence of distinct axons that output the integrated signal from the dendrites, such computations may be performed in a local compartmentalized manner along the neurites (**Figure 1b)**. In this case, local activity may be non-linearly integrated in a compartmentalized manner and be transmitted to post-synaptic neurites from within the compartment and without evoking current changes across the entire neuron. Clearly, compartmentalized activity requires that synaptic contacts will be sufficiently close to one another as greater distances will increase the probability that local currents will be shunted away or simply dissipate without producing local activity on the neurite.

*C. elegans* animals offer an appealing system to systematically study the possibilities for local compartmentalized computations within simply-structured neurites. The wiring diagram of the nervous system was mapped, giving the morphology of each neurite’s skeleton and the positions of the synapses along the neurites ^11,13–17^. Conceptual network-analysis approaches were applied to this unmatched detail-rich connectome to infer how function may arise from structure alone. These network-wide studies demonstrated that the overall modular architecture of the network can support various global functions such as sensory integration, sensory-motor convergence, and brain-wide coordination, where smaller circuit motifs contribute functional roles such as noise filtering, signal amplification and synchronized motor outputs ^13–29^. These functions were predicted based on the connectivity matrix alone, data that denote which neurons are chemically or electrically coupled, including the number of the connections they share. However, none of the above reports used the physical coordinates of the synapses to study the possibility that functional roles may be implemented within the neurites via local compartmentalized activities (**Figure 1b**).

Interestingly, local compartmentalized activities within neurites were experimentally observed in a few of the *C. elegans* neurons. For example, a reciprocal and compartmentalized activity was documented within the RIA interneurons to control the directionality of head bending, and within the RIS interneuron, activity in one segment induces locomotion stop while activity in another segment promotes reversals ^30–34^. But how prevalent are these discrete compartmentalized computations within the neural networkã

Here, we have systematically studied whether local compartmentalized computations may constitute a significant emerging property in the *C. elegans* neural network. We found that synapses tend to be clustered along neurites significantly more than randomly expected or than dictated by anatomical constraints. These organizations are evident in mutually-synapsing neurons and in triple-neuron circuits indicating that key local functions may be implemented along discrete segments of the neurites. Moreover, a new rule emerged, where the more common partners a pair of neurons shares, the more tightly clustered are their synapses on their mutual partners. This synaptic architecture may facilitate local signal integration and coordinated activation of functionally-related downstream neurons. Parallel localized computations along neurites can greatly enhance the computational power of compact nervous systems made of simply-structured neurons, thereby providing the evolutionary basis for the emergence of such capacities.

## Results

### Compiling the database of synaptic coordinates

To compile a dataset of synaptic coordinates, we used the connectome data provided by ^15^. These data contain the skeleton maps of most neurons as well as the position of the chemical and electrical synapses along the neurites. Here, we focus on the connectome formed by chemical synapses and use the term synapse to refer to these chemical synapses. The position of the chemical synapses consists of the coordinates of the pre-synaptic densities, while presumptive post-synaptic cells are identified by the proximity of their membrane to the site of the pre-synaptic structure. For a subset of the neurons, their cell body position is also provided.

We extracted the data for the adult hermaphrodite and compiled it into a data structure that allows easy retrieval of relevant parameters, including anatomical positions of the neurons together with their neurite extensions and the positions of their chemical synapses within the nerve ring (provided in supplementary file 1). In total, the data includes 181 neurons located in the animal head together with the 3,977 chemical synapses between them (**Figure 2a-b, and Methods**).

**Figure 2.**
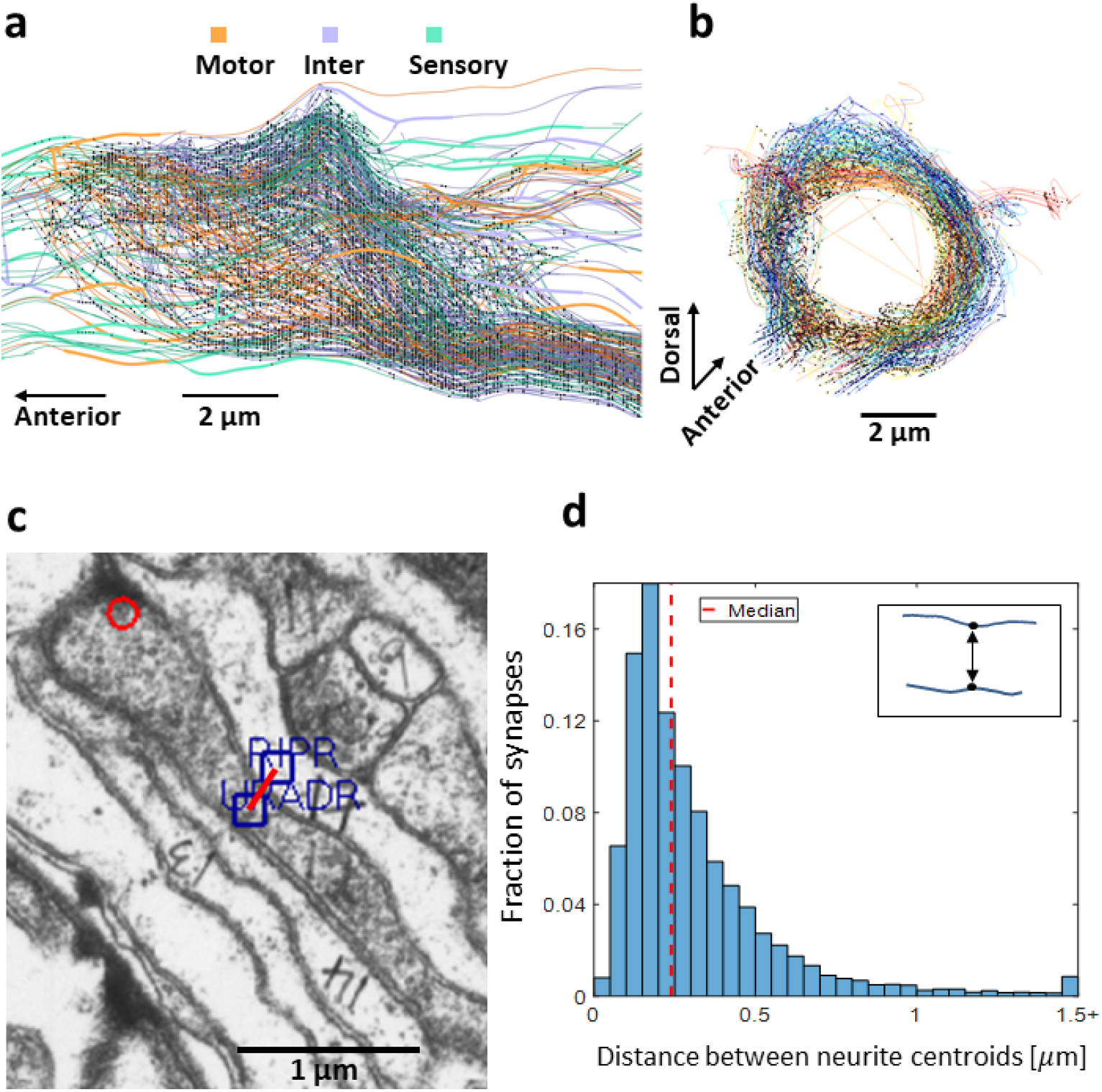
Compiling a database consisting of neurite’s morphology, synaptic positions along the neurites, and determining proximal neurite segments. **(a-b)** Anterior-posterior (a) and lateral (b) views of the neurites together with the chemical synapses in the nerve ring. Black dots mark chemical synapses. **(c)** An example of electron micrograph ^11^, reconstructed using the Elegance software ^35^, depicting the neurites of the presynaptic neuron URADR and its postsynaptic neuron RIPR. Blue rectangles mark the centroid of the neurites and the distance between the rectangles, denoted by the red line, is ∼0.24 µm. Red circle marks a presynaptic density, where URADR is presynaptic at a dyadic synapse to RIPR and a muscle arm from the head muscle dBWMR1 (synapse contin #5523). **(d)** Distribution of the distances between the centroid coordinates of the pre- and the post-synaptic neurites (see inset). Red dashed line marks the median of the distance distribution (=0.24 µm). We defined segments of two neurites to be proximate - where a functional chemical synapse could in principle be formed - if the distance between the neurites is shorter than this median value (see Methods).

To study the features of spatial synaptic distribution along neurites, it is mandatory to consider all the possible regions where functional synapses could in principle be formed. This can be done by determining which neurite segments are sufficiently close to one another to support such synapses (see example in **figure 2c**). However, the typical distance between neurite segments that share a synapse is unavailable. We therefore used the data above to compute the distance between neurites at the point that they form a synapse: For each presynaptic density, we retrieved the closest centroid coordinates of the pre- and post-synaptic neurites, and calculated the distance between them (**Figure 2c-d**). This provided the distribution of the distances between two neurites at the points where they form a synapse. The distribution of these distances is skewed such that the median distance is 0.24 μm, but for a significant number of synapses, the distance is greater than 0.5 μm (**Figure 2d**). Thus, to determine which neurite segments are sufficiently proximate to form a functional synapse, we conservatively set this threshold to be the median 0.24 μm (red dashed line in **Figure 2d**). As this threshold marks the distance between neurite centroids, the median synaptic distance is actually shorter. Hereafter, we term segments of two neurites to be proximate if their distance is within this median threshold.

Notably, we found that the length of proximate segments between two neurites provides a very good predictor for synapse existence, and the longer are the proximate segments, the higher the probability for these neurites to share a synapse (**Supplementary note and supplementary figure S1**).

### Synaptic contacts form clusters, significantly more than randomly expected

An elementary building block of a neural network consists of a chemical synapse that connects between one pre- and the one post-synaptic neuron. Often, such connected neurons share multiple synapses along their neurites. These synapses may be randomly distributed along proximate regions of the neurites, or alternatively, form clustered structures (**Figure 3a-b**). The latter case may hint to possible local compartmentalized functions.

**Figure 3.**
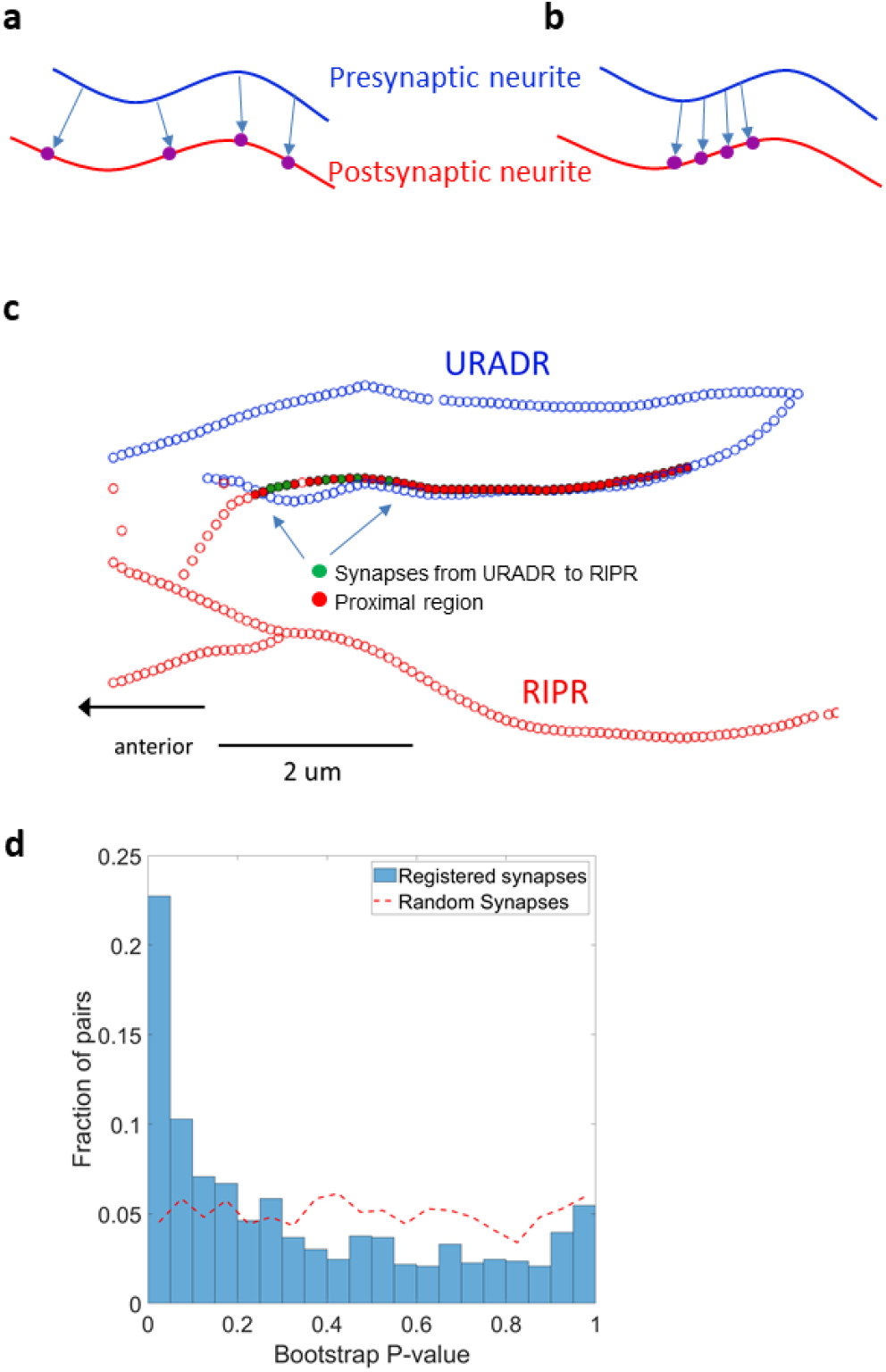
Synapses are more clustered than randomly expected based on anatomy alone. **(a-b)** The pre- and post-synaptic contacts can be either randomly (possibly uniformly) spread (a), or spatially clustered (b) along the neurite. **(c)** An example of two neurons sharing clustered synapses. The URADR neuron (blue) is presynaptic to the RIPR neuron (red). Empty circles mark the neurites, and the filled red circles show the proximate regions between the neurites where a functional synapse could in principle be formed (based on analysis shown in **Figure 2d**). The filled green circles denote the positions of the known registered synapses. Evidently, the synapses are clustered, rather than uniformly distributed, along the proximate regions of the neurites. **(d)** A histogram of the significance distribution of clustered synapses along all pairwise proximate neurites. P-values were calculated based on a bootstrap analysis on all pairs of neurons sharing two or more synapses along their proximate neurites (see Methods). For >30% of the neuron pairs, the mean distance between actual registered synapses is shorter than expected, and over 22% of the neuron pairs had a mean distance that was lower than that of 95% of the random instances drawn for a pair of neurons. As expected, a uniform-like distribution (dashed red line) is found when comparing between the randomly positioned synapses themselves.

To discern whether synapses form random or clustered distributions, and to assign a quantitative measure to their spatial organization along the neurites, it is essential to consider all the possible positions along the neurite in which synapses could in principle be formed. For this, we computed the distance between all pairwise pre-synaptic densities in all neurites, and constructed for each neuron a symmetric adjacency matrix for all pre-synaptic densities, where each entry *A*^*n*^_*ij*_ specifies the distance between a pair of pre-synaptic densities *(i,j)* along the neurite of neuron *n* (**Supplementary file 1**). We considered the distance along the neurite, rather than the Euclidean distance, as this measure reflects the extent of possible compartmentalized activity (**Supplementary figure S2**).

Analysis of this matrix demonstrated that the synapses are indeed clustered within proximate regions of the neurites in over 20% of the neural pairs (**Figure 3c-d**). To provide a quantitative significance for this clustering, we performed a bootstrap analysis: For each pair of neurons sharing two or more synapses, we compared the mean distance between the actual registered presynaptic densities and the mean distance between presynaptic densities that were randomly positioned along proximate regions of the neurites (red-circled segments in **Figure 3c**, and see Methods). This analysis revealed that synapses are clustered along proximate segments of the neurites, significantly more than randomly expected (mean distance between actual and randomly-positioned synapses was 3.1 μm and 4 μm, respectively, p<10^−13^, Wilcoxon rank sum test, n=500 pairs, **Figure 3d**). Thus, while synapses could in principle be randomly distributed along proximate segments of the neurites (while forming functional synapses), a significant fraction of them form clustered structures along the neurites.

### In- and out- going synapses between mutually-synapsing neurons form separate clustered groups

Mutually synapsing neurons represent a simple circuit embedded within neural networks (**Figure 4a**). A bidirectional connectivity may facilitate key computational roles, such as signal amplification if both synapses are excitatory ^36^, and a switch-like transition (an exclusive OR computation) in case of a reciprocal inhibition ^37,38^. We therefore asked if synaptic organization may support these functional roles locally along the neurites.

**Figure 4.**
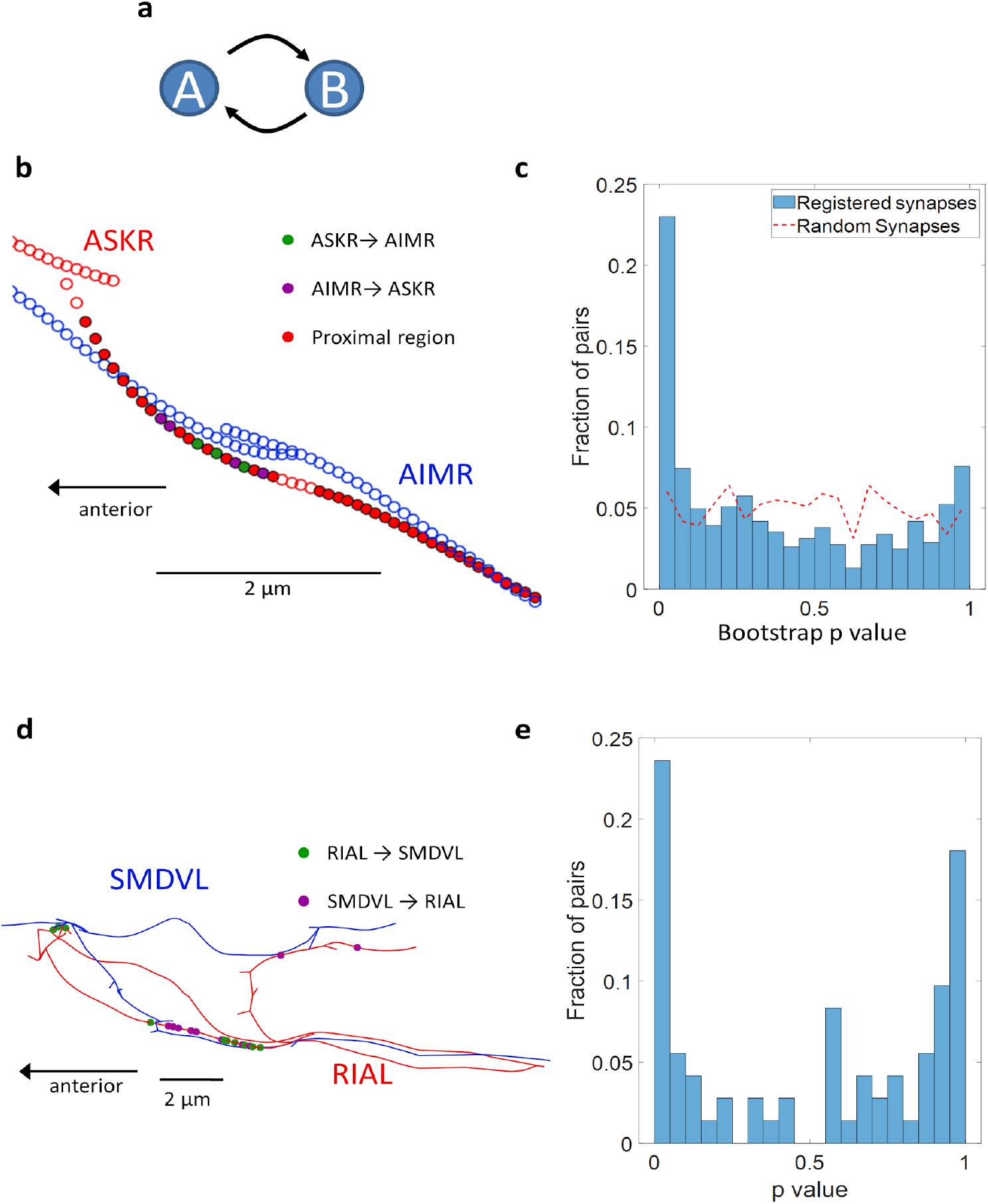
In- and out- going synapses between mutually-synapsing neurons form distinct clustered groups along proximate regions of the neurites. **(a)** Circuits comprising mutually-synapsing neurons may support key functional roles within the network. **(b)** An example of clustered synapses positioned along proximate neurite segments of the mutually-synapsing neurons ASKR and AIMR. Empty circles mark the neurites and red filled circles denote proximate regions on the neurites in which the distance allows synaptic formation (as defined in Figure 2d). Purple and green filled circles mark the locations of the actual registered synapses along the proximate neurites. **(c)** A histogram showcasing the distribution of the significance (p values) for the clustered synaptic organization between mutually synapsing neurons. In nearly 25% of these mutually synapsing neurons, the mean distance between actual registered synapses is significantly shorter than the expected distance obtained by randomly placing the synapses on proximate sections of the neurites (bootstrap p-value <0.05). Dashed red line marks a baseline distribution of the p values when comparing between randomly positioned synapses. **(d)** Within the synaptic clusters formed between mutually-synapsing neurons, incoming and outgoing synapses maintain distinct segregated clusters. Shown in an example for such segregated clustering in the mutually-synapsing neurons RIAL and SMDVL. The clustering is shown along the RIAL neurite. Note the segregation between the incoming synapses to RIAL (purple circles) and the outgoing synapses from RIAL (green circles). **(e)** Comparison of the distances between incoming and outgoing synapses shows that these form segregated clusters. For each pair of neurons that shared at least two synapses in each direction, we compared the distances between synapses with the same directionality with the distances between synapses with opposite directionality (see Methods). Presented is the distribution of p-values of one-sided ranksum comparison for all such pairs, indicating that synapses are more clustered by their directionality (incoming or outgoing) than expected by chance (Fisher’s combined probability test, p<10^−10^, n=72 pairs).

To study this possibility, we first analyzed whether bidirectional synapses are co-clustered within proximate regions. For this, we used the same approach as before, and calculated the mean distance between synapses in mutually synapsing neurons regardless of polarity. We found that the mean distance between registered synapses is significantly shorter than the mean distance between randomly-shuffled synapses found in mutually-synapsing neurons (3.7 μm and 4.3 μm respectively, p<10^−6^, Wilcoxon rank sum test, n=384 pairs, **Figure 4b-c**). Thus, just as for uni-directional connections between neurons, synaptic connections between mutually-synapsing neurons tend to be clustered along proximate regions of the neurites more than randomly expected.

Interestingly, within these synaptic clusters, incoming and outgoing synapses tend to cluster separately, such that the incoming synapses are segregated from the outgoing synapses (**Figure 4d**). To systematically quantify this segregated clustering, we analyzed pairs of neurons that share more than two synapses in each direction. For each such pair of neurons, we calculated the pairwise distances between all the synapses in one direction, and separately, for all the synapses in the opposite direction. We then compared these distances with the distances between oppositely directed synapses. These analyses showed that in over 20% of the bidirectional synapses (out of 72 in total), the distance between synapses going in the same direction is significantly shorter than the distances between opposite-directed synapses (p<0.05, bootstrap analysis, **Figure 4e**). Of note, similar segregated clusters are also found when comparing the positions of chemical synapses and electrical gap junctions (**Supplementary figure S3**).

### Tightly clustered synapses, connecting three-neuron circuits, may carry functional roles

We next considered the synaptic spatial organization in circuits made of three neurons

{A,B,C} that form simple connectivity layouts devoid of feedback or feedforward connections (**Figure 5a I-III**). In these circuits, the neurons {A and B} are either pre- or post-synaptic to a mutual neighbor neuron {C}. We therefore hypothesized that if the contacts of {A,B} are sufficiently close along neurite {C} then they may support local compartmentalized functional roles. For this analysis, we do not consider the locations of the proximal regions between A and C and between B and C, which might overlap or be statistically close along neurite C.

**Figure 5.**
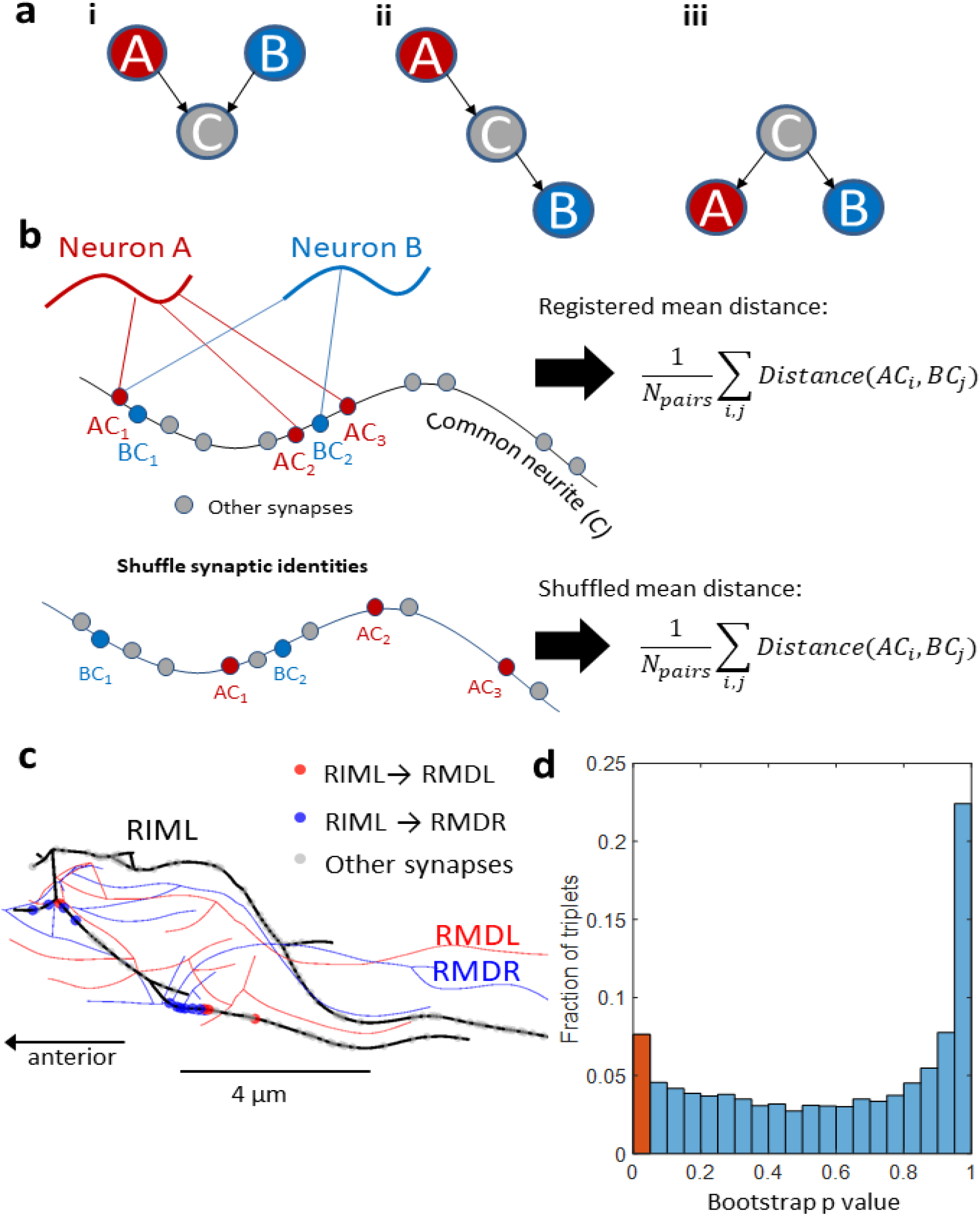
Three-neuron circuits, connected by clustered synapses, may support local functional roles. **(a)** Three possible layouts for a pair of neurons {A,B} that form synaptic connections with a third common neuron {C}: (i) the two neurons {A,B} are presynaptic to neuron {C}. Signals may be integrated locally on neurite {C}. (ii) Neuron {A} is presynaptic while neuron {B} is postsynaptic to neuron {C}. In this case, there is no computation per se, but the activity from {A} is transmitted to {B} locally on {C}, without involving the entire neuron {C}. (iii) Neuron {C} is presynaptic to both {A and B}, so that local activity on {C} can be simultaneously transmitted to both {A} and {B}. **(b)** To identify significant tightly-clustered synaptic contacts of {A,B} along neurite {C}, we calculated the mean distance between the known registered synapses (mean distance between red and blue dots, top), and compared it to the mean distances obtained after shuffling the positions of {A} or {B} synapses with the positions of synapses formed on neurite {C} by other neurons (bottom). A pair of synapses from {A,B} neurons on a common neuron {C} is considered to be significantly close if the mean distance of the registered synapses is lower than 95% of the shuffled synapses. **(c)** An example for a tightly-clustered set of synapses. Synapses from RIML to RMDR and RMDL are spatially clustered on the presynaptic RIML neurite. Note there are two such clusters. **(d)** The distribution of P-values following a bootstrap analysis for all the triple sets of neurons that we analyzed. Red bar marks the triplets that were defined as tightly clustered.

If {A and B} are both presynaptic to neuron {C}, then sufficiently close synaptic positions may lead to an (often non-linear) integrated activity in a local compartmentalized manner without spreading the signal along the entire neuron C (**Figure 5a, example I**). Another possibility is that activity of neuron {A} is transmitted to neuron {B} via local changes on neurite {C} without involving a global change in the entire neuron {C} (**Figure 5a, example II**). Finally, sufficiently close presynaptic positions on neurite {C} may simultaneously transmit similar activation patterns to both neurons {A and B}, a design that may support a timely-coordinated signaling (**Figure 5a, example III**).

To systematically identify such circuits, we considered all pairs of neurons {A,B} that form synapses with a third common neuron {C}, and computed the distances between the chemical synapses formed by the pairs of neurons {A,B} along neurite {C}. To provide a quantitative significance measure for how close (clustered) these synapses are along neurite {C}, we shuffled between the synapses formed by {A} or {B} neurons with other synapses formed by neuron {C} and other neurons (see Methods). We then compared the mean distance between the {A} and {B} synapses before and after the shuffle (**Figure 5b**). We defined the synapses of two neurons to be significantly clustered on the common neuron if the mean distance between them was shorter than that of 95% of the shuffled sets of both {A} and {B} synapses. An example of a triad (type iii in figure 5a) is shown in figure 5C: the synapses from RIML onto RMDL and RMDR form two clusters, where in each cluster, the synapses to RMDL and RMDR are tightly clustered.

We found that ∼8% of such triple-sets of neurons form synapses that are spatially significantly clustered (red bar in figure 5d and see supplementary file 2). For over 20% of the triplets, the synapses of the two neurons are actually positioned significantly farther away from one another, more than randomly expected (**Figure 5d**, rightmost bar with P value ∼ 1). This is because in most circuits, pairs of neurons will synapse on different sections of the common neurite and shuffling between synapses often yields a mean distance that is shorter than the genuine one. Nevertheless, the fact that nearly 10% of the synapses are significantly clustered suggests that these synaptic circuits may carry functional roles locally along the neurite.

We next studied the interesting triple-set neural circuits that form Feed Forward Loops (FFL). FFLs are recurrent motifs found in various biological networks, including the *C. elegans* neural network ^29^. In these circuits, one neuron {A} controls activity of a second neuron {B}, and both {A,B} control activity of a third neuron {C} (**Figure 6a**). The ubiquitous appearance of these circuits in biological networks suggests that they may carry key computational roles, including noise filtering, coincidence detection and more ^15,39–41^. In contrast, cyclic circuits form anti-motifs, and hence, are underrepresented in biological networks ^29^ **(Figure 6b)**.

**Figure 6.**
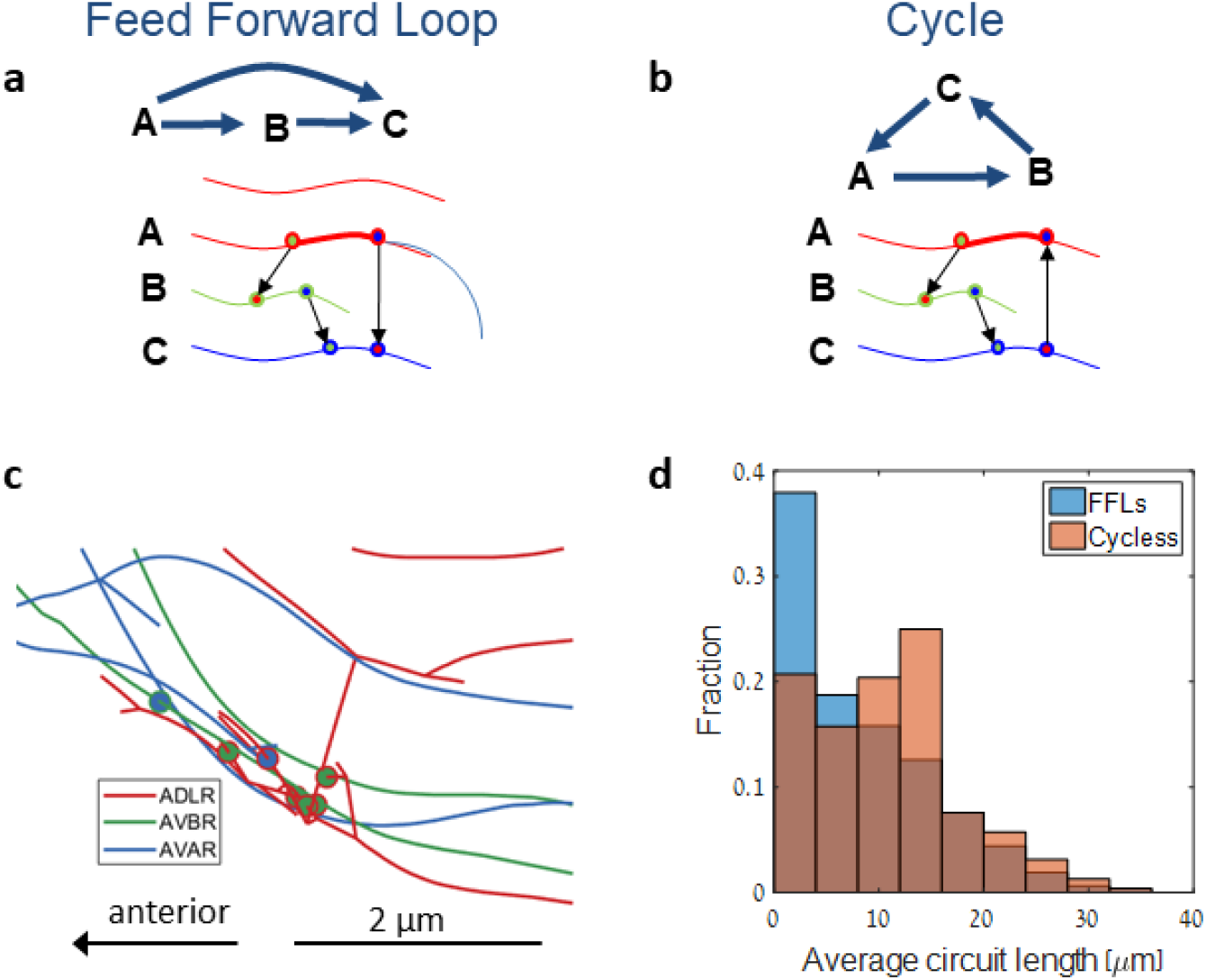
Feed Forward Loop functions may be implemented via local synaptic activity. **(a**,**b)** Network (top) and physical (bottom) schematics of a Feed Forward Loop (a) and a cycle (b). Thick line marks the largest distance between synapses of the circuit which we defined as the circuit length. **(c)** An example of a localized FFL circuit formed between the neuron ADLR, AWAR and AVBR. Circles mark synapses, where the inner color marks the synapse target neuron and the outer color marks the synapse origin. **(d)** A histogram showing the distribution of circuit lengths of FFLs and cyclic circuits. We considered all triple-set neurons that form an FFL or a cycle. For each synapse on neuron A, we measured the minimal circuit length (as shown in panels a and b). In cases of multiple A->B synapses, we averaged over all the circuits they form.

We therefore studied the intriguing possibility that FFL circuits may be hardwired via local synaptic connectivity along neurites, thus implementing their functional roles in a local compartmentalized manner. For this, we analyzed the positions of the synapses that form FFLs (and cycles) and calculated the shortest path along the neurites on which these circuits are implemented. The circuit length was defined as the longest neurite segment that connects two of the circuit’s synapses (bold red neurite in **Figure 6a-b**). Interestingly, we found that the mean circuit length for FFLs is significantly shorter than that of cycles (median of 5.7 μm in FFLs and 9 μm in cycles, n = 1096 and 79 respectively, p<10^−4^, Wilcoxon rank-sum test, **Figure 6c-d**).

Moreover, the mean number of synapses between neurons forming an FFL is higher than the number of synapses within cyclic circuits (**Supplementary figure S4)**. While this finding could partially explain why FFLs are characterized with synapses that are spatially closer to one another, their widespread occurrence (> 1,000 instances) suggests that localized FFLs may implement computational roles in a compartmentalized manner along neurites and without involving global changes of the entire neurons.

### Synaptic clusters follow the common neighbor rule

An extension to the simple three-neuron structure is known as the common neighbor rule. According to this rule, the probability for two neurons to be connected increases with the number of the mutual synaptic partners that they share ^18,42,43^. In analogy, we asked if the probability of two neurons to form close synapses on the neurite of their common target neuron increases the more common targets these two neurons share (**Figure 7a**). Analysis of the synaptic distance between triplets of neurons (as described above) revealed that indeed the probability for synapses of a pair of neurons to be clustered along their common neighbor neuron increases the more common neighbors they share (p<10^−5^, **Figure 7b**). Furthermore, the more common neighbors a pair of neurons shares, their synapses are positioned significantly closer to one another along the neurite of their common neighbor (p<10^−10^, **Figure 7c**).

**Figure 7.**
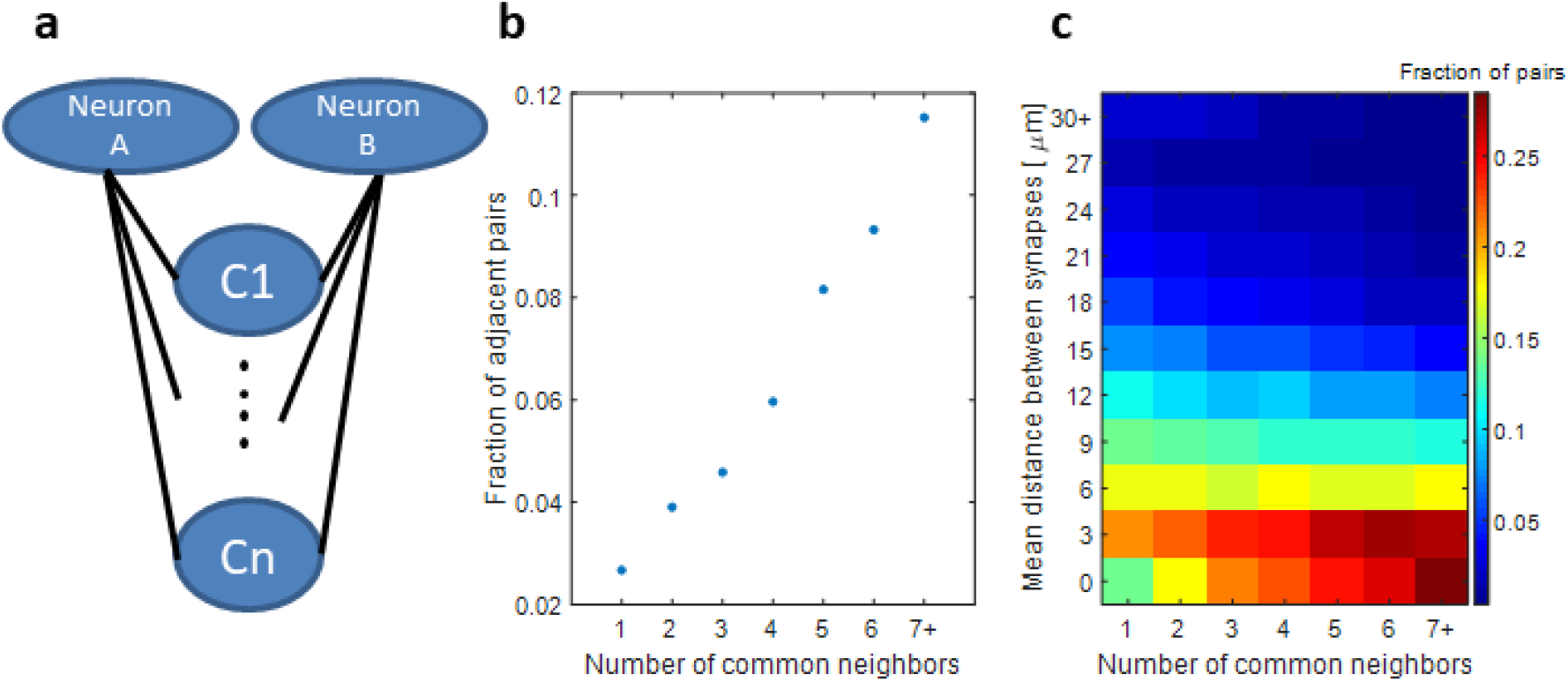
The more common neighbors a pair of neurons shares, the closer are their synapses along the common neuron. **(a)** Schematics showing a pair of neurons A and B together with their common neighbors C_1…n_. **(b)** Fraction of pairs of neurons, whose synaptic contacts on the common neighbor neuron are tightly-clustered, increases with the number of common neighbors that they share (definition of tightly-clustered synapses as in **figure 5**). **(c)** The mean distance between pairs of synapses decreases with the number of common neighbors that their neurons share. Shown is a histogram of the mean distance between the synapses of a pair of neurons along a common neurite, conditioned by the number of close neighbors that they share. The sum of each column is normalized to 1.

These findings may be somewhat expected as a pair of neurons sharing many common neighbors may synapse these neighbors within a restricted region of the nerve ring. Nonetheless, such anatomy may also carry functional roles as neurons that have many common neighbors are also more likely to be functionally related. Interestingly, the lateral left-right symmetry pairs of neurons, which often fulfill the same function, have significantly more common partners (p<10^−11^, Wilcoxon rank-sum test). Moreover, although these neurons typically reside on opposite symmetric sides of the nerve ring, their synapses are more likely to be closely spaced on their mutual partner than the synapses of two other random neurons (13% of the right/left pairs are clustered vs 8% of all others, p<10^−7, 2^ test). These findings indicate that synapses sharing the same functional roles will tend to be tightly-clustered on their mutual target neurons, further underscoring the possibility that local compartmentalized activity is an evolved feature in the network.

### Specific neural classes within structural motifs tend to form statistically close synapses

Next, we asked whether specific neural classes form and interact in the tightly-clustered synapses positioned along neurites. For this, we considered the well-defined classification of the layers in the *C. elegans* neural network, consisting of sensory-, inter-, and motorneurons. Neurons from each of these three classes (layers) can synapse neurons from other classes, including neurons from the same class (**Figure 8a**). We focused on the simple connectivity layouts analyzed for triple-set neurons above **(Figure 5a)**, this time keeping record of the class identity for each of the three neurons (**Figure 8a-b**).

**Figure 8.**
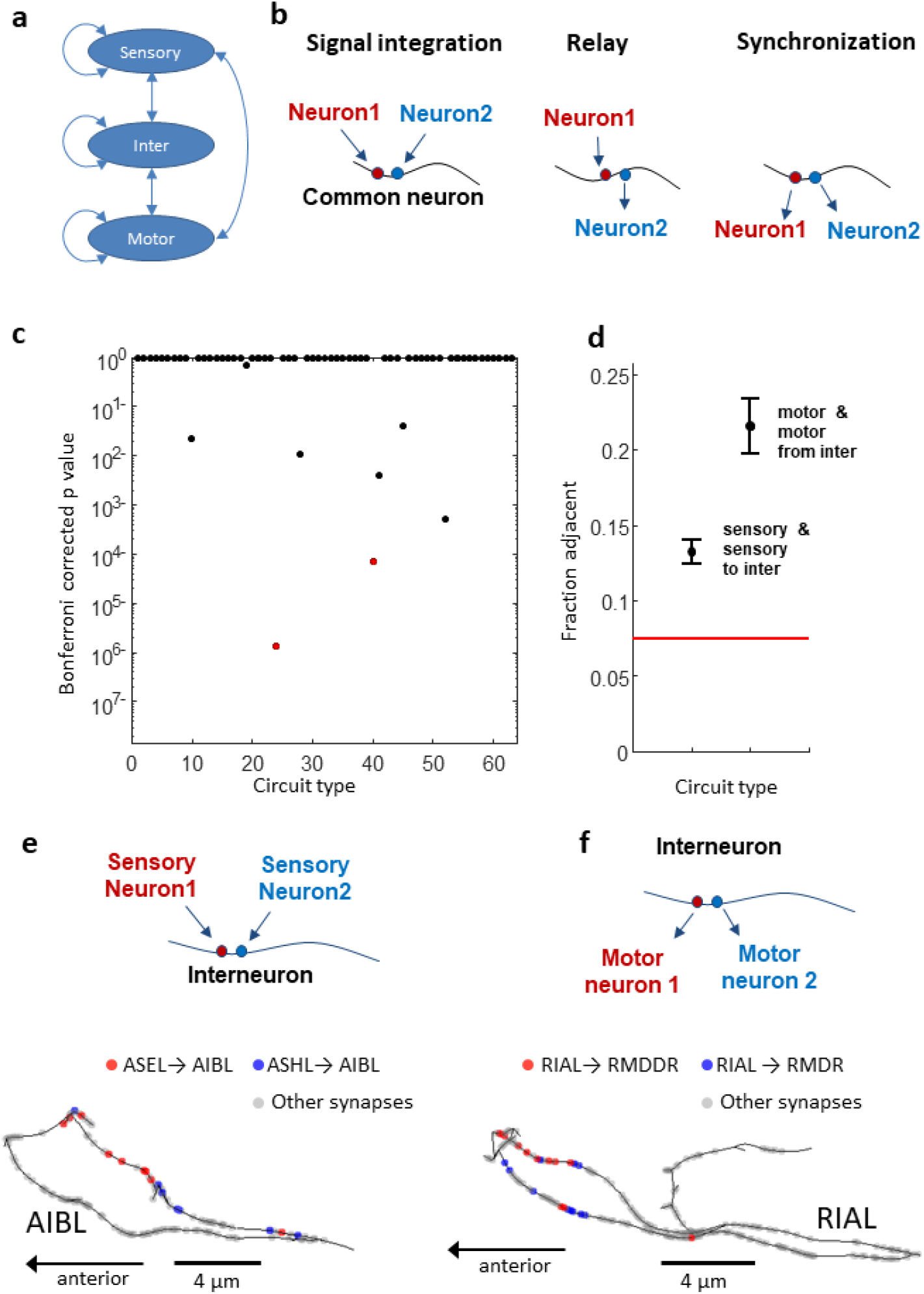
Synapses of specific neural classes are more likely to be clustered to one another. **(a)** Each of the *C. elegans* neuron classes (sensory-, inter-, and motorneurons) form synapses within and across the different classes. **(b)** Simple triple-set neural topologies may support localized functional roles: integration of signals, signal relay, and synchronized activation of two downstream neurons. **(c)** Of the 63 possible circuit types, only few are significantly overrepresented as forming clustered synaptic structures. A logistic regression analysis was used to determine statistical significance. P-values are following a bonferroni correction (see Methods). **(d)** The fraction of the two most significant motifs that form clustered synaptic contacts (out of the total significant motifs): {sensory1+sensory2 -> interneuron}, a circuit that may function as signal integrator; {interneuron-> motor1+motor2}, a topology that supports coordinated and synchronized activation. Red horizontal line denotes the fraction of motifs with such clustered synapses across all triplets. Error bars denote the standard error of the mean assuming a Binomial distribution. **(e-f)** Illustrations of the two motifs together with an actual example. **(e)** Top: Two presynaptic sensory neurons that converge on a postsynaptic interneuron. Bottom: the sensory neurons ASHL and ASEL form clustered synapses on the proximate region of their mutual postsynaptic interneuron AIBL. **(f)** Top: A presynaptic interneuron synapses two motor neurons. Bottom: The interneuron RIAL is presynaptic to two motor neurons, RMDR and RMDDR. The two synapses on the proximate region of RIAL are closely positioned. Note all the other synaptic connections that these interneurons form are marked in grey

For three different layouts, where each of the three neurons can be either sensory, inter- or motorneuron, there are 63 possible circuit combinations. Of these 63 combinations, few circuits emerged as forming clustered synaptic connections, significantly more than randomly expected (**Figure 8c-d**, and see Methods for statistical considerations). The two most significant circuits were particularly intriguing: (i) Two sensory neurons forming clustered post synaptic contacts on a third common neurite (layout i in **figure 8b**); (ii) An interneuron that is presynaptic to a pair of motor neurons, and whose presynaptic contacts are tightly-clustered together (layout III in **figure 8b**). Notably, these two motifs consist of a pair of neurons belonging to the same neural layer (either sensory or motor) and with same synaptic directionality (either pre- or post-synaptic) to the common neuron. Moreover, in both motifs, the common neuron is an interneuron, suggesting that local compartmentalized computations may be more prevalent in interneurons than in other neuronal classes.

These circuit motifs point to interesting functional roles: for sensory neurons, this motif may support integration, or coincidence detection, of environmental stimuli performed on the common neurite of the interneuron. For example, the two chemosensory neurons ASEL and ASHL form clustered synapses along their common postsynaptic interneuron AIBL (**Figure 8e)**. Responses of each of these sensory neurons to external cues may be locally integrated on the AIBL neurite without necessarily involving activity changes in the entire neuron. A motif where two tightly-clustered presynaptic sites form post-synaptic contacts on two motor neurons may support fine coordinated muscle activity. For example, the RIAL interneuron synapses two motor neurons, RMDR and RMDDR, which innervate head muscles controlling head position and movement directionality (**Figure 8f**).

## Discussion

Quantitative analyses of connectomes typically abstract the connectivity data into an adjacency matrix, where each neuron is treated as a single homogenous node in the network ^14,15,26–29^. Such approaches ignore valuable information regarding the physical location of the synapses, which in the case of *C. elegans*, is available. By combining adjacency matrices together with the position of the synapses along the neurites, we revealed that synaptic distribution along neurites is not random, nor can it be explained by mere anatomical constraints. Instead, the fine clustered synaptic organization hints to a compartmentalized activity that may provide key functional roles.

For example, clustered post synaptic contacts can support local nonlinear integration of individual inputs, and reciprocal clusters between mutually-synapsing neurons may support various functions in a local compartmentalized manner along the neurite and without involving activity changes in the entire neuron (**Figures 3-4**). Overall, this synaptic clustering phenomenon appears in a substantial fraction (∼30%) of the proximal neurites (**Figure 3d** and **figure 4c,e**). Similarly, we demonstrated that clustered organization of synapses is found predominantly in specific types of tri-neuron circuits, further underscoring the high prevalence for evolved, rather than random, synaptic organization that may fulfill synaptic functional roles (**Figures 5-8**).

We delineated a new synaptic rule, analogous to the known ‘common neighbor rule’ found in neural networks ^18,43^. According to the ‘common neighbor rule’, the more common partners a pair of neurons shares, the higher the probability for the pair of neurons to be connected. We found that the more common neural partners a pair of neurons shares, the more statistically close together are the synapses of the pair of neurons on their mutual partners (**Figure 7**). As sets of neurons forming such circuit connectivity typically function together, the tightly-closed position of the synapses along the neurites can facilitate relevant functional roles in a local compartmentalized manner.

Moreover, specific neural types, forming unique structural motifs, are enriched within tightly-clustered synapses: Sensory neurons are more likely to form localized post-synaptic contacts on interneuron neurites, and presynaptic contacts on interneurons are more likely to be clustered together when they synapse downstream motoneurons (**Figure 8**). These layouts are particularly enriched in the network and may support intriguing functional roles in a synaptic local manner. Activities of two sensory neurons, whose postsynaptic contacts are closely positioned on a mutual postsynaptic neurite, may be integrated locally. In this case, the information of two different environmental cues, each sensed by a different sensory neuron, may be integrated and processed locally within the interneuron neurite. Similarly, when two motor neurons receive inputs from the same neurite region of the presynaptic interneurons, the local activity in the interneuron can be further transmitted in an efficiently correlated manner. This is particularly useful for motor outputs which require fine coordinated activity between various motoneurons.

These findings point out that interneurons are the primary neurons that implement local compartmentalized activity along their neurites (**Figure 8**). Indeed, several studies already demonstrated compartmentalized neural activity in the *C. elegans* interneurons ^30,32–34^. A well-studied example is the RIA interneurons, which control the animals’ head bending due to compartmentalized and reciprocal activity between the dorsal and the ventral parts of its neurite. In our analyses, we detected clustered synapses outgoing from the dorsal compartment of RIA to the dorsal motor neurons (e.g., RMDR and RMDDR, **figure 8f**). As worms crawl along their dorso-ventral axis, such spatially-clustered connectivity may support the worms’ gradually-enhanced head bending, a locomotive behavior that underlies the weathervane chemotaxis strategy ^32–34,44^.

Compartmentalized activity was also observed in the RIS interneuron, in the region between the nerve ring and the axonal branch ^30^. The RIS cell body is posterior to the nerve ring and its neurite extends anteriorly, encircling the nerve ring. Posterior to the nerve ring, the synaptic neurite branches to form an axonal branch. Interestingly, the nerve ring segment and the axonal branch show compartmentalized activity, where activity in the nerve ring process alone correlates with locomotion stop, while co-activation together with the axonal branch promotes reversal events ^30^. This secluded local activity along the axon/neurite nicely exemplifies the dual role of this interneuron in dictating stops and reversals, presumably to allow sensory integration and behavioral outputs, respectively.

To determine proximal segments of two neurites, we used a conservative estimation calculated to be lower than the median of the distances between neurites (**Figure 2c-d**). The use of this conservative threshold suggests that the actual number of synaptic clusters is actually higher, and hence, the number of neurites with possible compartmentalized computations is also higher than predicted herein. Furthermore, the majority of the synaptic clusters are found along a single continuous segment of the proximal neurites, which in many cases also constitutes a significant fraction of the neurite (see for example, **figures 2c, 3c, 4b**). This indicates that the observed synaptic clusters are not due to possible fragmented segments of the proximal neurites, further underscoring their significant non-random clustering that may support functional roles.

When studying tri-node circuits, we considered the positions of the synapses and did not separately define the proximal regions. Synapse clustering in these cases may result from proximity of the proximal regions. When analyzing these types of triple-set circuits, we find that specific circuit types (*e*.*g*. sensory-sensory-interneurons, **figure 8**) are significantly overrepresented as having a marked short distance between their synapses. This suggests that local activity may be particularly relevant in these types of tri-node circuits.

Dyadic and triadic synapses, where a single presynaptic density makes synaptic contacts with two or three different neurons, respectively, are abundant in the *C. elegans* nervous system (66% of all synapses ^15^). In the nerve ring alone, 61% of all synapses share more than one post-synaptic partner. As the distance between the position of the presynaptic contacts of dy/tri-adic synapses is zero, such structures could contribute to the observed clustered synaptic formation and the emerging motif layouts along the neurites. Nevertheless, the mere existence of such structures may actually further underscore the directed evolution to form such clusters, which presumably carry fine functional roles along the neurites. For example, a dyadic structure formed between an interneuron and two postsynaptic motor neurons may better support a synchronized output that is essential for coordinated undulatory locomotion of the animal.

Interestingly, the anatomically distant right and left pairs of neurons tend to form clustered postsynaptic connections, further strengthening the possibility that synaptic organization may be sculpted to support compartmentalized computation along neurites. These bilateral neurons are often considered to be functionally equivalent, and are likely to respond to the same neurodevelopmental signals that guide them to their postsynaptic partners. If this signal is localized at a particular site in the postsynaptic neurite, then the neurons responding to that signal will form their presynaptic densities opposite to that site. This developmental process may explain synaptic clustering of functionally-related neurons. However, whether this clustering is meant to support local computational roles, or is a consequence of other constraints (e.g. economy in neural wiring ^45,46^) requires further experimental evidence. Of course, these two possibilities are not mutually exclusive. Our computational analyses provide a comprehensive list of neurons and circuits, including the distance between the associated synapses (**Supplementary files 1-3**), thus offering candidate circuits for studying compartmentalized activity via functional imaging experiments.

Another intriguing possibility for the emergence of local compartmentalized activity is the support of experience-dependent plasticity. Classical experiments demonstrated that local synaptic activity marks (‘captures’) the synapse and targets it with proteins that support long-term plasticity in a process known as synaptic tagging and capture ^47,48^. As *C. elegans* animals can form various memory types, ranging from simple non-associative habituation to classical conditioned associative memories ^49–52^, local synaptic activity can efficiently mark specific target synapses. Such selected tagging increases the memory capacity of the network as neurons can be partitioned into distinct synaptic modules, each undergoing plasticity processes independently of the others.

Taken together, local compartmentalized activities, facilitated by the clustered synaptic organizations revealed herein, can enhance computational and memory capacities of a neural network. Such enhancement may be particularly relevant for animals with a compact neural network and with limited computational powers, thereby explaining the evolutionary forces for the emergence of these synaptic organizations.

## Methods

### Dataset used in this study

The data used in this study was extracted from ^15^, and is provided in https://wormwiring.org/. The structure of the database is detailed in ^35^. Herein, we used the ‘contin’, ‘display2’ and ‘synapsecombined’ tables of the N2U dataset. The ‘contin’ table provides an individual identifier, the contin number, of each neuron. The ‘display2’ table contains the 3D structure of each neuron and the coordinates of the neurons’ synapses. Here we focused on the chemical synapses, which we refer to throughout simply as synapses. The ‘synapsecombined’ table provides the pre- and post-synaptic partners of each chemical synapse. As our focus was the structural properties within the nerve ring, we considered the 20-microns of the worms’ anterior part, removing neurites and synapses that are posterior to that position.

### Formatting the dataset

The dataset ^15^ (https://wormwiring.org/) provides the coordinates of the pre-synaptic densities as well as the centroids of the pre- and the post- synaptic neurites. The post-synaptic cells were identified by their proximity to the membrane where the presynaptic density is located. Thus, we positioned the postsynaptic structure on the closest centroid coordinate of the postsynaptic neurite.

Each synapse in the dataset is registered on the presynaptic neuron and on each of the postsynaptic neurons. For synapses that were not registered on all partners, an attempt was made to add the registration on the missing neurons by locating the neurite segment closest to the synapse location. In cases where the closest segment was more than 2 µm away from the synapse, the synapse between the two neurons was removed.

In cases of dyadic to triadic synapses, where each presynaptic neuron has several postsynaptic partners, we registered each such synapse separately for each postsynaptic partner. All these copied registrations of the synapse had the same position as that of the original synapse (so that the distance between them is 0).

In addition, the structure of few neurons in this dataset appears fragmented and disconnected, probably due to missing data in the original EM images or segmentation errors. For these neurons, we filled the gaps by ‘stitching’ disjoined ends. For disjoint segments harboring short gaps, where the intact connectivity was apparent from the known neurite anatomy, we manually stitched the segments. When such ‘stitching’ was not possible, small disconnected parts of the neurite were removed from the main neurite.

### Quantifying significance of synaptic clusters between pairs of neurons

To provide a quantitative measure for synaptic proximity, formed between two neurites, we applied the following procedure: First, we marked the coordinates of proximate neurites (less than 0.24 µm, a threshold calculated based on **Figure. 2d**) on a pair of neurons. These coordinates denote the positions in which synapses could be formed theoretically based on distance alone (red circles in **Figure 3c**). We then compiled a set of possible synaptic coordinates that include the actual registered synaptic positions and the proximate neurite segments. Next, the synapses between the two neurons were randomly placed in the set of possible synaptic positions, and the mean distance between them was calculated. This process was repeated 200 times to generate the distribution of mean distances between randomly placed synapses. We then calculated the fraction of the randomly placed instances that have a smaller mean distance than the real set of synaptic locations. This fraction was calculated for each pair of neurons in which the presynaptic neuron shares more than one synapse with the postsynaptic neuron (**Figure 3d**), as well as for a pair of mutually-synapsing neurons (**Figure 4c**).

### Evaluating prediction of synaptic connectivity between pairs of neurons based on cell-body positions or length of proximate neuritis

To evaluate the difference in predictability of synaptic connectivity between pairs of neurons, we used a logistic regression to predict if a pair of neurons is connected based on either the distance between their cell bodies or the length of close neurites that they share (as seen in **Supplementary figure S1a,c**). We then compared the area under the curve of the receiver operating characteristics (AUC-ROC) of the two models^42^. Indeed, we find that the AUC is higher when predicting based on neurites’ close sections (AUC of 0.85) than when using the distance between cell bodies (AUC of 0.61). We used a non-parametric test for the difference between the AUCs^53^ to find that this difference is indeed significant(p<10^−5^).

### Quantifying synapses pairwise clustering

To evaluate the clustering of incoming and outgoing synapses between pairs of neurons, we focused on pairs of neurons that share more than two synapses in each direction. We then grouped the distances between pairs of synapses into two groups: within group distances (distance between synapses that are in the same direction) and between groups distances (distance between synapses of opposite directions). Finally, we used the one-sided ranksum test to check whether the between-groups distances are significantly longer than the within-groups distances. The results are shown in **Figure 4e**. The same method was also used to evaluate clustering of chemical and electrical synapses (**Supplementary figure S3b**).

### Evaluating clustered synaptic connections between neurons on a common neurite

For each triplet of neurons {A,B,C} in which both A and B share chemical synapses with C, we consider three possible types of connectivity: (A→C, B→C) ; (A→C, B←C) ; (A←C,B←C). For each of these connections, we compared the mean distance between the A and B synapses along the C neurite to the distance we would randomly expect. To approximate the random distance, we randomly chose synapses from all the synapses on neuron C and position either the A or B synapses in these locations (**Figure 5a-b**). We used this method to generate 1000 shuffles of the A synapses and 1000 shuffles of the B synapses. A pair of neurons was defined as ‘tightly clustered’ on a common third neuron if the registered mean distance between A and B synapses was smaller than 95% of both A and B shuffles. When the total of possible shuffled instances was less than 1000, we considered all possible instances.

### Logistic regression for tightly-clustered motifs

To examine the effect of specific motifs on the probability that two neurons will form tightly-clustered synapses on a third neuron, we used logistic regression. As we wished to test the specific effect of the triplet type, we accounted for the following mediators by assigning them features in the regression: the number of synapses each neuron had on the common neighbor (two features), the number of common neighbors that the pair of neurons had (one feature), the type of the common neuron (two binary features) and the direction of the synapses each neuron had on the common neuron (two binary features). To these common features, we added the binary feature ‘isTriplet’, which was changed in each regression in accordance with the motif examined. In total, 63 different regressions (one for every possible motif) were constructed, each trying to predict which neural triplets will have clustered synapses. Shown in **Figure 7c** are the bonferroni corrected P values (corrected for 63 comparisons) as derived from the regression to the hypothesis that the weight assigned to the ‘isTriplet’ feature is greater than 0.

### Analysis of FFL and cycle circuits

For each FFL or cycle circuit that we detected in our dataset, we considered each of the synapses from the first neuron of the motif to the second one (A→B synapses in **Figure 6a-b**). For each such synapse, we measured three distances that define the motif: (1) the distance along the neurite of neuron A to the closest A-C synapse. (2) The distance between the A-B synapse and the closest B-C synapse along the neurite of neuron B. (3) The distance between the A-C synapse and the B-C synapse along the neurite of neuron C. The circuit length was defined as the longest of these three distances. Repeating this process for each A-B synapse allowed us to compute the average circuit length of the triplet. The distribution of these average lengths is shown in **Figure 6d**.

#### Supplementary file 1

A matlab file containing the network data used for all the analyses described herein. It is an array of structures, one for each neuron. Each structure contains the following fields:

**NameStr** - the neuron name.

**type** - the neuron type (sensory neuron / interneuron / motoneuron).

**Contin** - the contin number assigned to the neuron according to ^15^.

**CellBodyX / Y / Z** - the 3D coordinates of the cell body center of mass, if annotated

**uniqueObjs** - a table with the object elements that form the neuron. Each object has an ID, a 3D coordinate, and links to other objects to form the neurite skeleton.

**adj_mat** - a matrix whose entries describe the distances along the neurite between the neuron objects. A_ij_ is the distance along the neurite between elements i and j in the uniqueObjs table.

**all_synapses -** a table that describes all the synapses on the neuron’s neurite with the following fields:

<u>Idx:</u> unique synapse ID, that can be used to detect the same synapse on the different neurites it connects.

<u>Post1-4:</u> marks the postsynaptic targets of each synapse. In cases where a single synapse on the neuron connects to more than one postsynaptic target, each target is registered separately in the table, with the post1 field indicating each separate target while the idx field remains identical.

<u>sections:</u> The number of EM sections in which the synapse appears (an approximation to the synapse size).

<u>x,y,z:</u> the 3D coordinates of the synapse registration on the neurite.

<u>isOriginal:</u> Marks whether the synapse registration on the neurite was created to account for discrepancies in the dataset or existed in the original data (see data formatting in methods).

<u>pre/postobj:</u> The objectID of the synapse pre and postsynaptic elements. adjMatInd: The synapse row number in the adj_mat table of the neuron.

**Dist_mat -** a matrix whose entries describe the distances between the synapses along the neurite. A_ij_ is the distance between elements i and j in the all_synapses table.

#### Supplementary file 2

contains a csv table with the results used for detecting tightly-clustered synapses of pairs of neurons on a common neuron (shown in **Figures 5-8**). The table contains the following fields:

**commonNeuron** - the common neurite.

**N1/2name** - Names of the two compared neurons.

**synapseType1/2** - the direction of the chemical synapses examined, with respect to the common neuron (toNeuron for example means synapses that are directed to the common neuron).

**probShuffleA/B** - the computed probability that the shuffling process (see methods) of the synapses of each neuron will result in a shorter mean distance between the synapses of the two neurons than the original mean distance.

**meanDist** - the mean distance between the synapses of neurons 1 and 2.

**meanShuffledDist** - the mean distance between the two neurons over all shuffles

**nSynapses** - the number of synapses of each neuron with the common neuron

**nPerms** - the number of possible synaptic shuffles.

**commonPartnersNum** - the number of common neighbors between neurons 1 and 2.

### Supplementary file 3

A list of the top common neurons and the number of significant triplets that they form.

## Acknowledgements

The Zaslaver group was supported by ERC (336803), ISF (1300/17), ICORE (1902/12). R.R was also supported by the Jerusalem Brain Center. S.W.E. was supported by NIMH grant R01MH112689. A.Z. is the Greenfield chair in Neurobiology.

## Data availability

All the data and the analyses scripts are publicly available through: https://github.com/zaslab/Neurite-computations

## Supporting online information

Supplementary note 2

Supplementary figures 3-6

Supplementary references 7

## Supplementary note

### Neurites’ proximity predicts synaptic connectivity between neurons

To support the suitability of our approach to analyze proximal segments of neurites as possible positions for synapse formation, we analyzed how well this approach predicts neural connectivity within the dense structure of the nerve ring. Specifically, we compared this predictive approach with the findings showcasing that the position of cell bodies may predict neural connectivity. The latter relies on the notion that in *C. elegans*, neuronal cell bodies are spatially positioned such that the total neurites’ length is minimized, so that connected neurons will have the shortest neurites needed to support synaptic connections ^1,2^. We considered the 77 neurons that innervate the nerve ring and for which the cell body position was available, and calculated the pairwise distance between the centroids of their cell bodies and sorted these pairs by the distance. We then divided the sorted list into groups of 100 pairs, and for each group, calculated the fraction of pairs that are synaptically connected as well as the mean number of synapses between all the pairs in the group (**Supplementary figure S1a-b**).

In agreement with the minimal neurite length principle, we found that the probability for two neurons to form a synapse increases the shorter the distance between their cell bodies (r= -0.79, p<10^−6^). However, the slope of this relation is very mild, and only 20% of the pairs of neurons whose cell bodies are less than 10 μm apart share a synapse (**Supplementary figure S1a**). These findings indicate that predicting the existence of a synapse based on the distance between two cell bodies is not readily feasible.

A much better predictor for the possibility to find a synapse is the length of proximate segments between two neurites (as defined in **Figure 2d**, p<0.001, see methods for calculating the significance). In this case, the probability to find a synapse between two neurons strongly correlates with the length of proximate sections that these two neurons share, where over 50% of the neurites with proximate segments longer than 4 μm are connected by one or more chemical synapses (**Supplementary figure S1c**). Furthermore, the number of synapses between the neurons also strongly correlates with the mean length of the proximate region (**Supplementary figure S1d**). Thus, neurites that tend to be proximate over great lengths are likely to share a synapse.

## Supplementary figures

**Supplementary figure S1.**
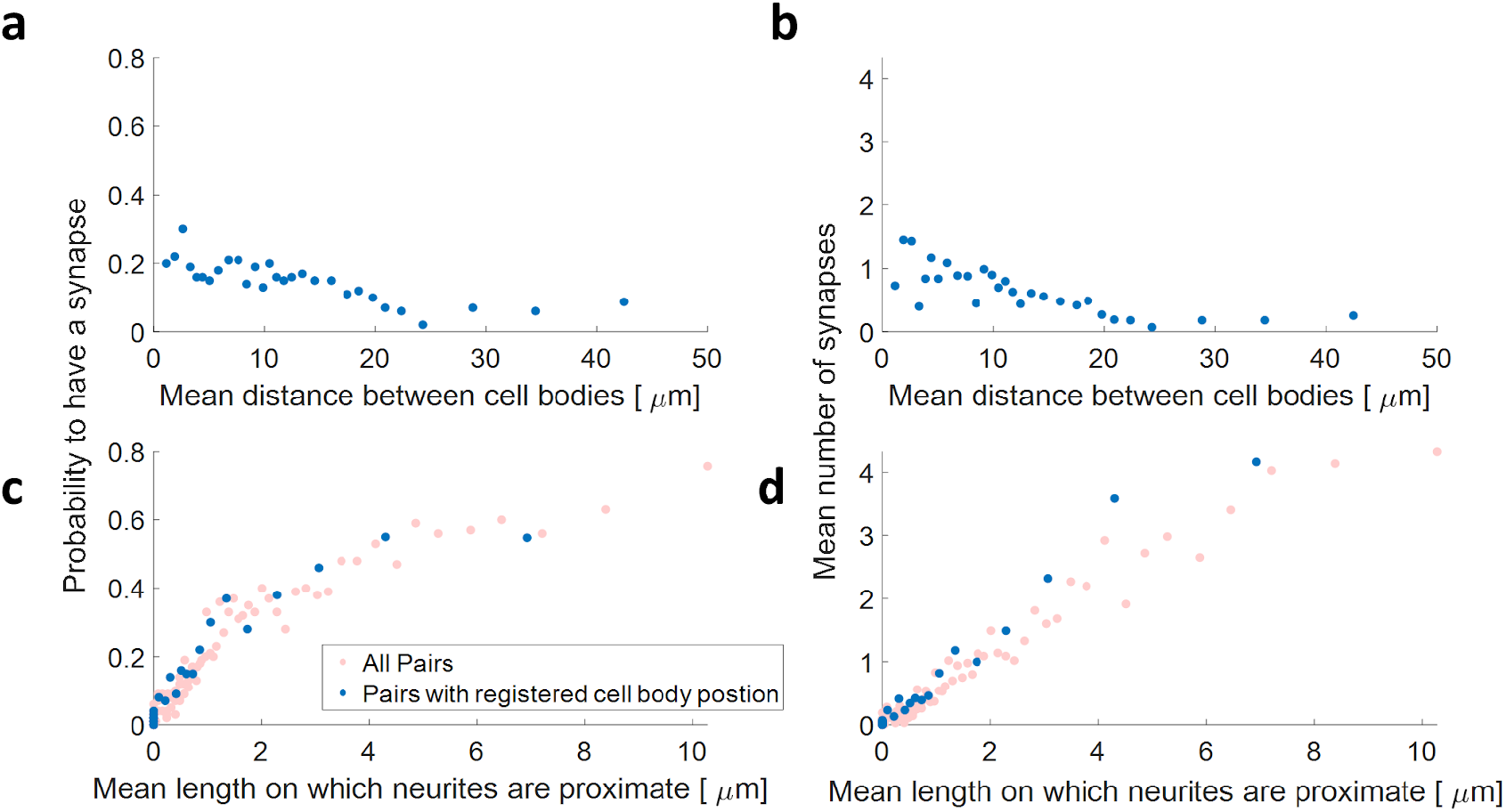
The length of proximate regions between neurites predicts synaptic connectivity.

(**a-d)** The fraction of pairs of neurons that share a synapse (a,c) and the mean number of synapses between neurons (b,d) are plotted as a function of the distance between the neuronal cell bodies (a,b) or the length of proximate neurite segments, along which a synapse can be formed. Proximate neurites are defined as having a distance not greater than 0.24 µm (see Figure 1c). Each dot is the mean of 100 pairs of neurons. Blue dots show pairs for which the cell body coordinates are available (registered) in the dataset. Note that neurites’ proximity is a much better predictor for the existence of a shared synapse (p<0.001, see methods).

**Supplementary figure S2.**
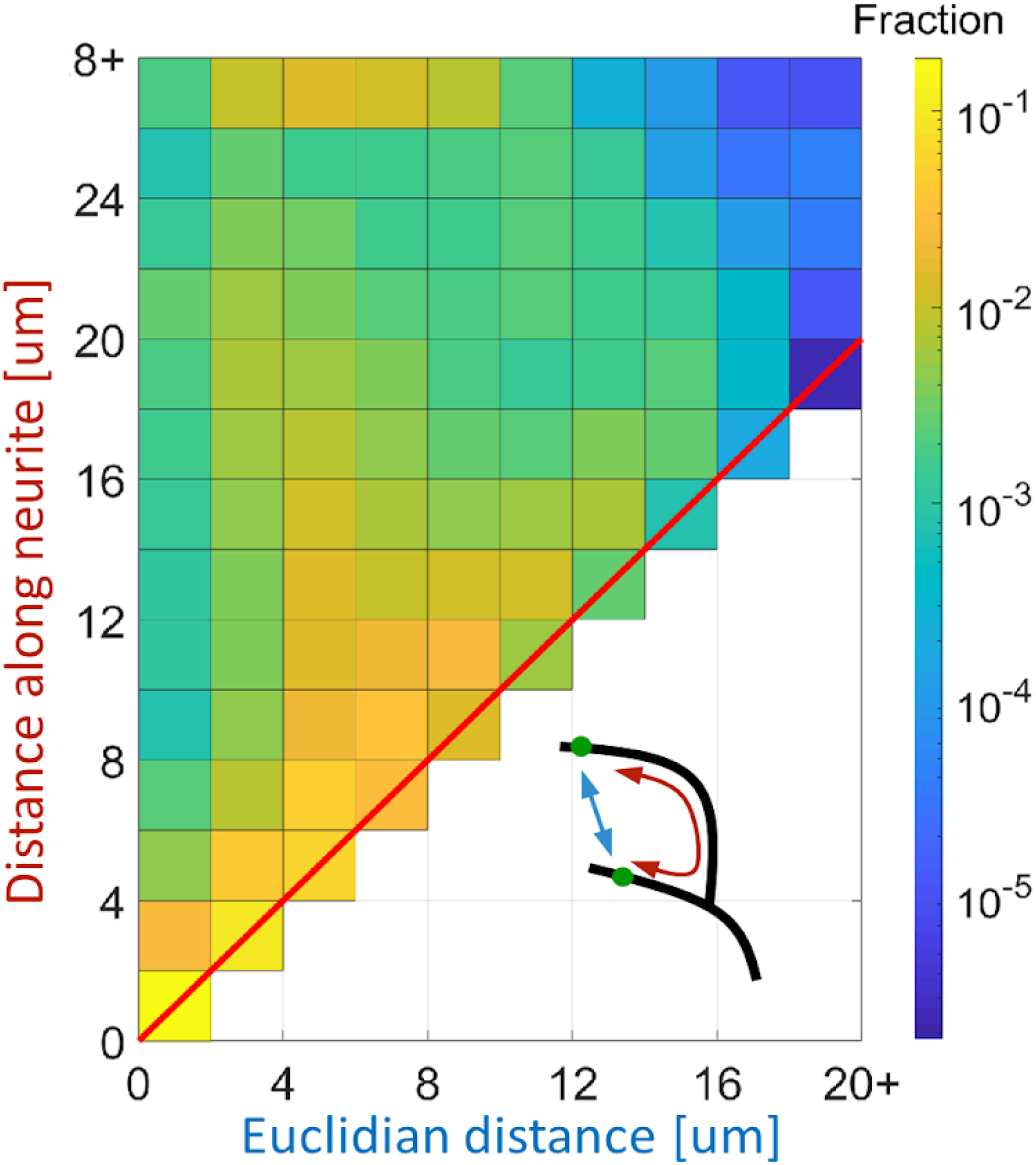
The distance between synapses along the neurite is significantly longer than the euclidean distance. We measured the Euclidean distance (blue) and the distance along the neurite (red) between all pairs of pre-synaptic densities (as depicted in the bottom right diagram). Shown is the joint histogram of these two distances for all pairs. As seen, for many pairs, the distance along the neurite is 2-3 times longer than the euclidean distance due to neurites curvative anatomy. Color marks density distribution in log scale. Red line marks the diagonal.

**Supplementary figure S3.**
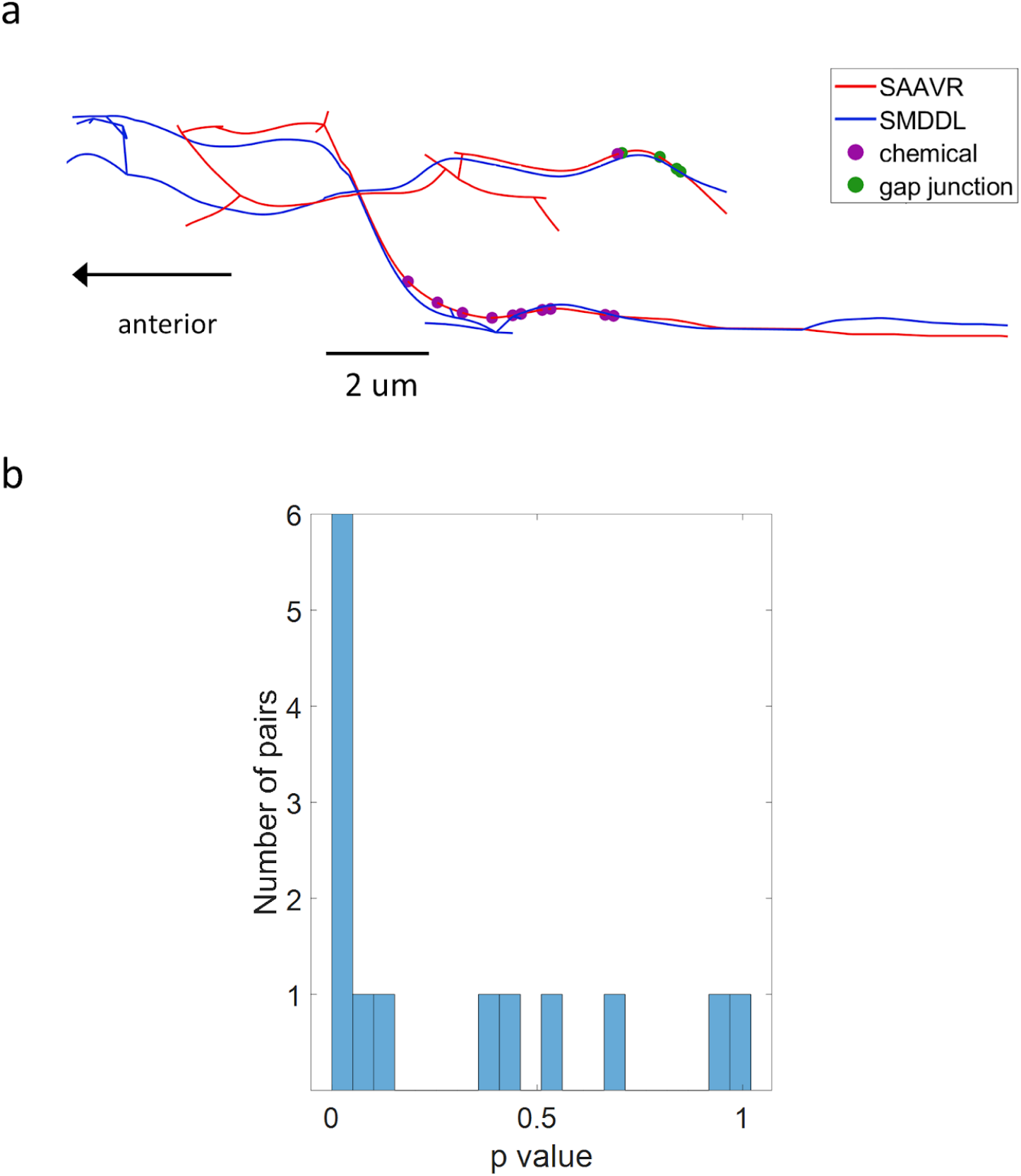
Chemical synapses tend to cluster separately from electrical synapses.

(a) An example for separate clusters for chemical and electrical synapses between the SAAVR and the SMDDL neurons. Notice the separation between chemical and electrical synapses.

(b) Connectome-wide evaluation of separation between chemical and electrical synapses. For each pair of neurons that share two or more synapses of each type, we compared the in-group distances with the between-groups distances of the synapses. Shown is the distribution of p-values of a one-sided ranksum comparison for all such pairs. From analyzing the distribution of these p-values, we find that more synapses are clustered by their type than expected by chance (Fisher’s combined probability test, p=6*10^−5^, n=14 pairs).

**Supplementary figure S4.**
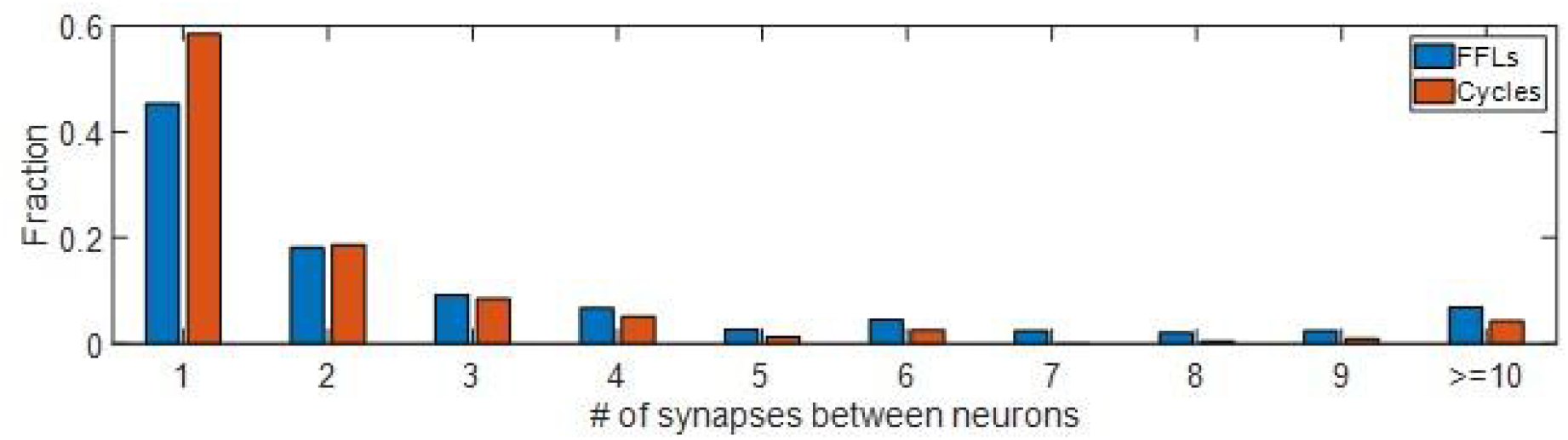
The connections of Neurons in FFL circuits have more synapses than in cycles. The distribution of the number of synapses between the different neurons that participate in FFLs and cycles. On average, FFL circuits have more synapses than cycles (p<10^−5^, Wilcoxon rank-sum test, n=3288 and 237 pairs of neurons, means of 3.3 and 2.3 synapses, respectively).

